# Microsaccades track location-based object rehearsal in visual working memory

**DOI:** 10.1101/2023.03.21.533618

**Authors:** Eelke de Vries, Freek van Ede

## Abstract

Besides controlling eye movements, the brain’s oculomotor system has been implicated in the control of covert spatial attention and the rehearsal of spatial information in working memory. We investigated whether the oculomotor system also contributes to rehearsing visual objects in working memory when object location is never asked about. To address this, we tracked the incidental use of locations for mnemonic rehearsal via directional biases in microsaccades while participants maintained two visual objects (coloured oriented gratings) in working memory. By varying the stimulus configuration (horizontal, diagonal, and vertical) at encoding, we could quantify whether microsaccades were more aligned with the configurational axis of the memory contents, as opposed to the orthogonal axis. Experiment 1 revealed that microsaccades continued to be biased along the axis of the memory content several seconds into the working-memory delay. In Experiment 2, we confirmed that this directional microsaccade bias was specific to memory demands, ruling out lingering effects from passive and attentive encoding of the same visual objects in the same configurations. Thus, by studying microsaccade directions, we uncover oculomotor-driven rehearsal of visual objects in working memory through their associated locations.

**SIGNIFICANCE STATEMENT:** How humans rehearse information in working memory is a foundational question in psychology and neuroscience. To provide insight into the cognitive and neural bases of working-memory rehearsal, we turned to microsaccades – small eye-movements produced by the brain’s oculomotor system. We reveal how microsaccades track the locations of visual objects during memory rehearsal, even when object locations are never asked about. This brings three advances. From a psychology standpoint, it demonstrates how memory rehearsal automatically engages object locations. From a neuroscience standpoint, it demonstrates how such location-based rehearsal relies on brain circuitry that also controls our eyes. Finally, from a practical standpoint, it demonstrates how microsaccades can be utilised to track the properties of working-memory rehearsal across space and time.

## INTRODUCTION

The brain’s oculomotor system, which controls our eye movements, has also been linked to the regulation of covert cognitive processes. To date, the cognitive involvement of the oculomotor system has been established most clearly in the studies of covert visual-spatial attention and the rehearsal of spatial information in working memory (e.g., 1–9). In the current study, we tested whether the oculomotor system also contributes to the location-based rehearsal of multiple visual objects in working memory when object locations are never explicitly asked about. To address this, we here studied microsaccades as a direct time-resolved output from the oculomotor system.

Studies in which participants are explicitly tasked with memorising the locations of visual stimuli have revealed that the oculomotor system plays a role in the mnemonic rehearsal of these locations. Most directly, spatial rehearsal of visual locations has been shown to trigger eye movements to these locations (10, 11). Complementary evidence comes from studies showing better visual-spatial memory when eye movements are allowed throughout the delay interval (12–16, but see also 17, 18), and worse memory when eye movements are prompted to other locations (16, 19–21). Neuroscience studies in humans (22–25) and non-human primates (26, 27) have further revealed how spatial locations in working memory are represented in the frontal eye fields – a key cortical node in the brain’s oculomotor system (e.g., 28, 29). These studies collectively point to an important role for the oculomotor system in the rehearsal of spatial-location information in working memory.

Besides the retention of locations, working memory enables us to retain object-specific visual features, such as object colour and shape. The aforementioned oculomotor involvement in visual working memory may be specific to the rehearsal of locations (as suggested, for example, in 12, 16). At the same time, we know from ample prior studies that space remains a profound organizing principle for object retention in working memory, even when the location of memoranda is never asked about. For example, spatial locations may help to keep memory objects separate, and facilitate the binding of features belonging to the same memory object (30–35). Likewise, spatial locations may act as a medium for object selection and prioristisation from working memory (36–39), and may serve ‘spontaneous’ object rehearsal in working memory (40). It has remained unclear, however, whether location-based rehearsal of visual object features other than location relies on the brain’s oculomotor system.

The aim of the current study was to address whether the oculomotor system contributes to working memory through location-based rehearsal of multiple visual objects, even when object location is never asked about. To address this, we capitalised on directional biases in microsaccades (cf. 2, 41, 42) as a direct and time-resolved read-out of oculomotor engagement during working-memory retention of two visual objects. By presenting memory objects in different spatial configurations at encoding (**Fig 1a**), we were able to isolate and track object rehearsal – as reflected in microsaccades – through the incidental locations of the memoranda. Furthermore, by comparing directional microsaccade biases across different task demands with the same spatial configurations (**Fig 1b-d**), we could disentangle the contribution of working-memory rehearsal from passive and attentive encoding of the same visual objects in the same configurations. Doing so, we reveal oculomotor rehearsal of visual objects in working memory through their associated – but in our case always task-irrelevant – locations.

**Figure 1.**
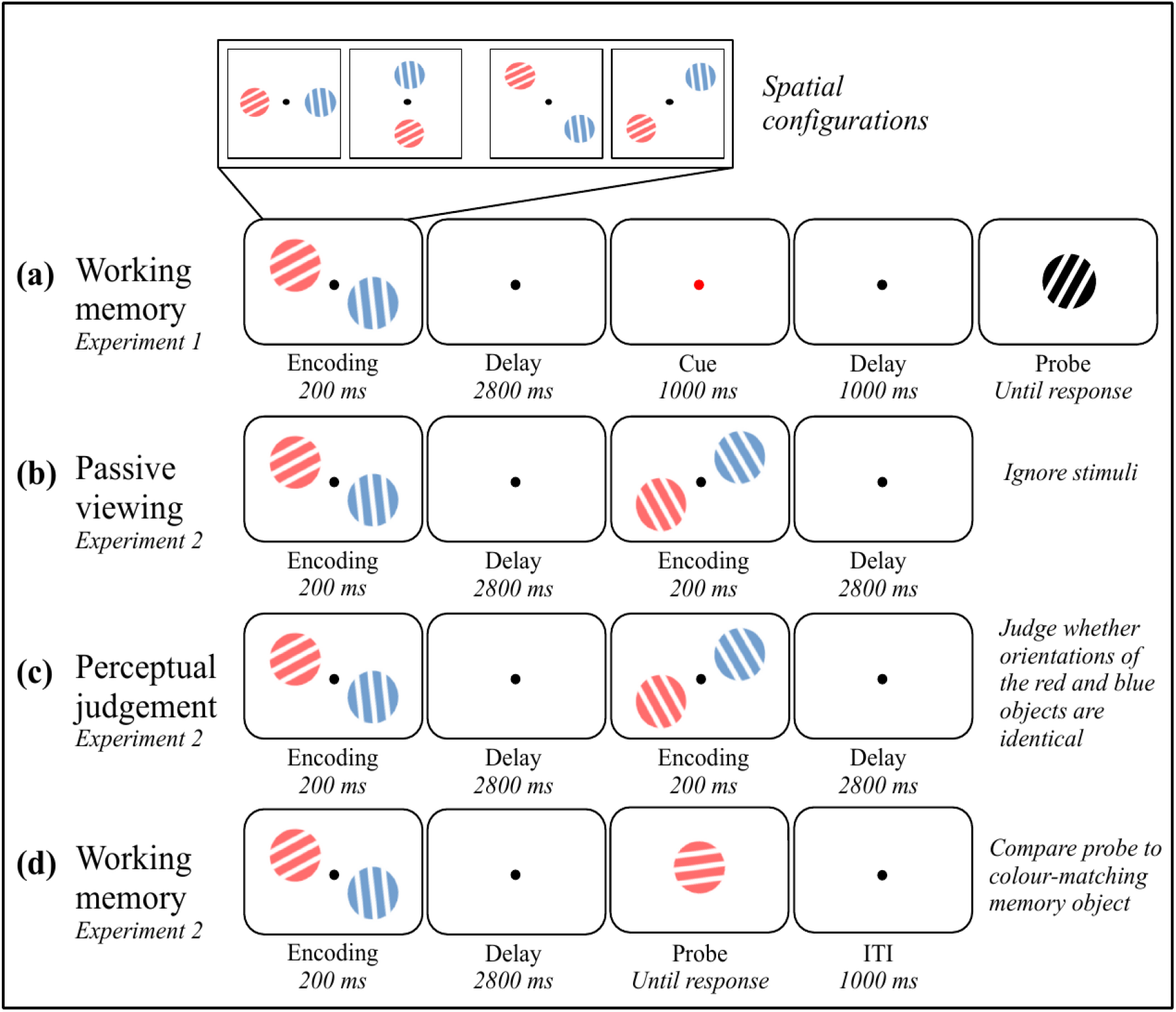
Task schematics of Experiment 1 (panel a) and Experiment 2 (panels b-d). In every case, trials started with the bilateral presentation of two sample gratings, followed by a delay interval. **a)** In Experiment 1, participants indicated whether a probe was oriented clockwise or counterclockwise with respect to the target memory object that was indicated by a colour retrocue presented during the delay. In Experiment 1, the two visual objects could occupy one of four configurations at encoding (horizontal, vertical, and either of two diagonal configurations). **b)** In the passive viewing condition of Experiment 2, participants were instructed to ignore the stimuli, while maintaining central fixation. **c)** In the perceptual-judgement condition of Experiment 2, participants had to indicate whether the orientations of the two simultaneously presented gratings in a given display were identical. **d)** The task in the memory condition of Experiment 2 was the same as in the first experiment, only now the cue was integrated into the probe, by virtue of the probe’s colour. In Experiment 2, the two visual objects always occupied either of the two diagonal configurations at encoding.

## RESULTS

### Experiment 1: Microsaccades reveal location-based oculomotor rehearsal in visual working memory

Before turning to our main eye-tracking results, we ascertained that participants were able to perform the working-memory task: participants had an average accuracy of 75.97% (*SD* = 11.37%) and an average reaction time of 1019 milliseconds (*SD* = 346 milliseconds).

The main purpose of Experiment 1 (**Fig. 1a**) was to investigate whether the oculomotor system is involved in the maintenance of visual objects in working memory, even when object location is never asked about. The incidental use of location for working memory was tracked by classifying saccades as “aligned” when their direction was closer to the configurational axis of the visual objects at encoding, and “misaligned” when their direction was closer to the orthogonal axis.

**Figure 2b** depicts the distribution of saccade sizes across the full 2.8-second delay period for trials with cardinal and diagonal stimulus configurations. The majority of saccades in the delay period were less than 1 degree visual angle (i.e. microsaccades, 41, 43–47) and did not revisit the initially encoded locations of the memory objects (which were presented at a distance of 6.4 degrees).

**Figure 2.**
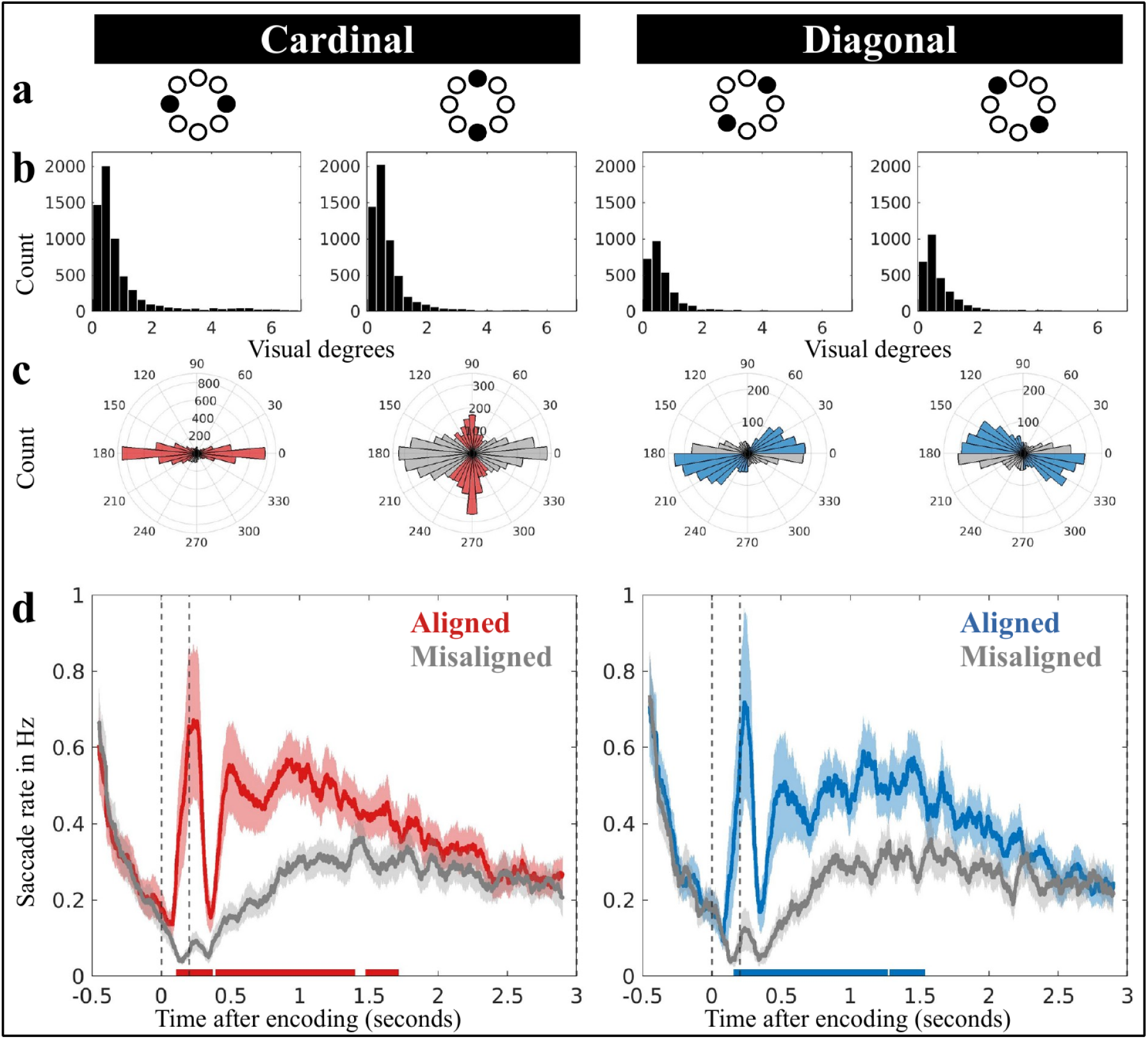
A directional microsaccade bias during working-memory maintenance reveals location-based object rehearsal. **a)** Spatial lay-out of the four stimulus configurations that were used in Experiment 1, with two solid black circles marking the locations of each stimulus pair at encoding **b)** Saccade-size distributions of delay-period saccades reveal that eye movements did not revisit the encoding locations of the memoranda. **c)** Polar distributions of delay-period saccades, with the coloured bins corresponding to saccades along the aligned axis, and the grey bins corresponding to saccades along the misaligned axis. **d)** The time courses of the saccade rates (in Hz) for saccades along the aligned and misaligned axes. The horizontal lines at the bottom highlight statistically significant clusters. Shaded lines represent ±1 standard error of the mean. The two vertical dotted lines mark stimulus onset and stimulus offset.

Our main result concerns the direction of the identified saccades as a function of memory-object configurations at encoding. **Figure 2c** depicts the polar distribution of all saccades detected during the delay period. The coloured bins correspond to saccades along the aligned axis, while the grey bins correspond to saccades along the misaligned axis. Taking all of the saccades during the memory delay into account reveals how saccade directions were generally biased along the axis of the visual objects at encoding (i.e. the aligned axis). While there is generally a horizontal bias in cardinal configurations (Fig. 2c, left panels; consistent with e.g., 2, 48), we found a configural memory bias on top of this general preference for horizontal saccades. This memory-configuration-specific bias is especially evident when comparing the two diagonal configurations, where the data is less distorted by the background horizontal bias (**Fig. 2c**, right panels).

To quantify this bias and track its evolution over time, we collapsed across horizontal/vertical and across the two diagonal configurations and counted saccades along aligned versus misaligned axes (relative to encoding configurations) as a function of time. Crucially, directions that were classified as aligned in one configuration would be classified as misaligned in the complementary configuration. Accordingly, by collapsing configurations, we could isolate condition-specific saccadic biases by subtracting out any ‘background’ saccade-direction preferences that would be common to all configurations. Further, note how the probe was always presented centrally, so any directional-microsaccade bias could not reflect probe anticipation.

**Figure 2d** shows the saccade rates across time for the aligned (coloured) and misaligned (grey) axes. A cluster-based permutation comparison of the time courses confirmed significantly higher saccade rates on the aligned than on the misaligned axis (indicated by the horizontal lines in **Fig. 2d**, cluster *P’s* < 0.001, 0.01, and 0.012 for the cardinal-configuration data; cluster *P’s* < 0.001 and 0.003 for the diagonal-configuration data). Importantly, this was not only evident during encoding, but lasted for a considerable period of time well into the working-memory delay interval.

Thus, these data show that during working-memory maintenance of visual objects, saccades – particularly microsaccades – occur more frequently along the spatial axis where memoranda were incidentally encoded than along the orthogonal axis (where no stimuli were encoded). This is consistent with object rehearsal by the oculomotor system through location, even when object location is never asked about.

### Experiment 2: The microsaccade bias during the delay reflects working-memory demands

In Experiment 1, microsaccades continued to be biased along the aligned axis well into working-memory delay. However, merely demonstrating that microsaccades remained biased in their direction during the delay does not necessarily imply that they reflect active memory maintenance, as such biases could also reflect residual, lingering consequences from prior sensory encoding.

Experiment 2 (**Fig. 1b-d**) sought to replicate the microsaccade bias during the memory interval (this time using only the two diagonal conditions; **Fig 3a**) and to compare it to two control conditions in which participants either (1) passively viewed the same objects, or (2) were required to make a perceptual judgement on the two objects (detect instances where both simultaneously presented objects had identical orientations) without subsequent working-memory demands. If the bias truly reflects active working-memory maintenance (rehearsal) processes, it should only be profound in our memory task, and not after passive presentation, or perceptual evaluation of the same objects, that we presented in the same configurations, followed by the same delay.

**Figure 3.**
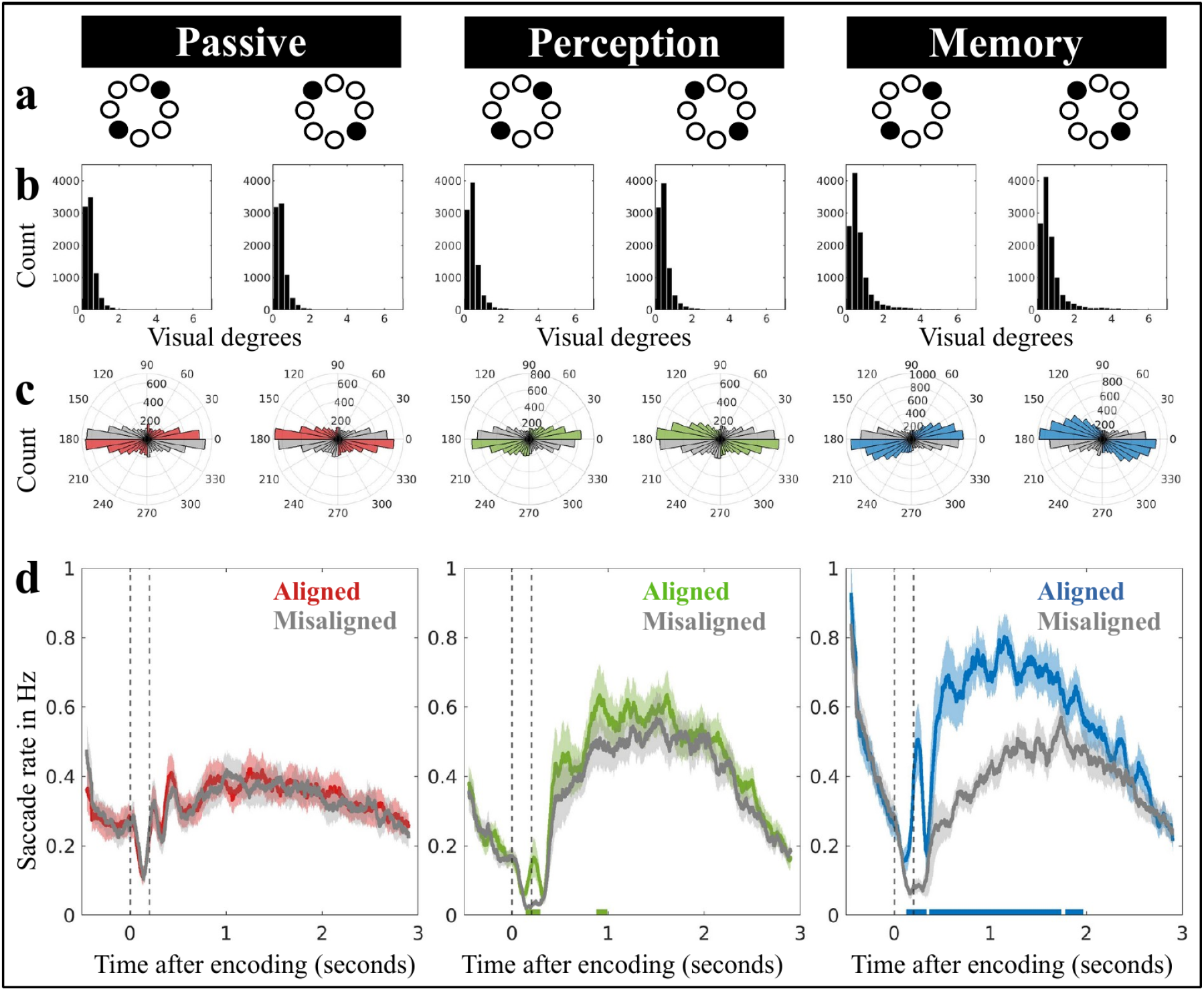
Directional bias in microsaccades is driven by working-memory demands. Spatial lay-out of the stimulus configurations that were used in Experiment 2, with the locations of each stimulus pair indicated by the two solid black circles. **b)** Saccade-size distributions of delay-period saccades. **c)** Polar distributions of delay-period saccades, with coloured bins representing saccades along the aligned axis and grey bins representing saccades along the misaligned axis. **d)** The time courses of the saccade rates (in Hz) for saccades along the aligned and misaligned axes. The horizontal lines at the bottom highlight clusters that are statistically significant. Shaded lines represent ±1 standard error of the mean. The two vertical dashed lines denote the onset and offset of the stimulus.

Before turning to our main eye-tracking results, we first confirmed that participants comprehended the working-memory and the perceptual-judgement tasks. In working memory blocks, average accuracy was 70.57% (*SD* = 13.32%) and average reaction time was 1348 milliseconds (*SD* = 616 milliseconds). In perceptual-judgement blocks, participants showed high hit rates following target displays (*M* = 83.38%, *SD* = 17.53%) and low false-alarm rates following non-targets displays (*M* = 2.60%, *SD* = 2.62%).

Having established that participants were able to do both tasks, we turn to our main eye-tracking results. **Figure 3b** shows the distribution of saccade magnitudes for passive, perceptual, and memory trials. As in Experiment 1, the vast majority of saccades occurred within the microsaccade range and again this was the case for all conditions. **Figure 3c** depicts the polar distribution of saccades during the delay period, again showing a clear bias on memory trials (in blue, on the right), but little evidence of a similar bias for the two control conditions with passive object presentation (in red, left) or the perceptual judgement task (in green, middle).

To quantify the bias and track its temporal profile, we again counted saccades along the aligned and misaligned axes, as defined by the incidental stimulus configurations at encoding. **Figure 3d** shows that saccades in passive blocks are equally likely to occur along the aligned and misaligned axes (cluster *P* > 0.3), indicating that stimulus presentation alone cannot account for the reported directional saccade bias in working-memory trials. In perception blocks, we discovered a slight bias for saccades to occur more along the aligned axis (cluster *P’s* = 0.01 and 0.019), but this bias was relatively weak and appeared to persist for only about 1 second after stimulus onset, which was also the time limit for participants to indicate that the two objects were identical.

In comparison, and replicating the results of Experiment 1, the memory condition again reveals a profound directional bias that persisted well into the working-memory delay period (cluster *P’s* < 0.001, 0.007, and 0.01). A direct comparison of the memory condition with both control conditions confirmed that the memory condition had a significantly stronger bias in microsaccade direction: the saccade rate effect (aligned minus misaligned saccades rates, averaged across the full a-priori defined delay period) was significantly larger in memory blocks (*M* = 0.19, *SD* = 0.11), than in passive blocks (*M* = 0.01, *SD* = 0.03), *t*(24) = 7.99, *p* < 0.001, and perception blocks (*M* = 0.05, *SD* = 0.06), *t*(24) = 6.07, *p* < 0.001.

These findings rule out that the observed saccade bias during working-memory maintenance is due to lingering processing of the passive viewing or attentive encoding of salient visual objects per se. Instead, the prolonged directional saccade bias in the memory task must be attributed to working-memory demands, which is consistent with an oculomotor-driven rehearsal of memorised visual object features (colour and orientation) through their memorised locations.

## DISCUSSION

It has been established that the brain’s oculomotor system not only controls where we look, but also participates in the rehearsal of spatial-location information in working memory (as reviewed in 4, 9–11, 14, 49). It had been called into question, however, whether this role extends to visual working memory of objects whose location is never asked about (see e.g., 12, 16). By analysing microsaccades as a direct and time-resolved output from the oculomotor system, we provide unique evidence that the brain’s oculomotor system also participates in the rehearsal of visual objects in working memory, even when object location is never asked about.

Experiment 1 revealed how microsaccade directions remained biased during the working-memory delay along the axis at which objects were incidentally presented at encoding. Experiment 2 replicated this central finding and ruled out lingering effects from passive or attentive encoding. By demonstrating directional biases in microsaccades during working-memory maintenance, we thus reveal rehearsal of memorised visual-object features through their incidentally encoded memorised locations – and implicate the oculomotor system in such rehearsal. In the following, we consider and discuss three central questions raised by these findings.

### What may the observed microsaccade bias during working-memory maintenance reflect?

Spatial biases in microsaccades have previously been linked to covert shifts of attention to peripheral locations (2, 41, 50, but also see 51, 52), including when directing attention within the spatial-layout of visual working memory (e.g., 42, 53, 54). Accordingly, our findings of directional microsaccade biases during working-memory maintenance may reflect active attentional rehearsal or refreshing of memory representations (55). Though the location was never asked about, it may still facilitate rehearsal. This is consistent with prior research demonstrating that space remains important for organising working-memory content, even when location is never asked about (e.g., 37, 38, 42, 53, 54). Our data go beyond this existing body of work by implicating the brain’s oculomotor system in such incidental use of location, and by demonstrating this role during working-memory maintenance (extending prior work implicating the oculomotor system in mnemonic selection; see e.g., 42, 53, 54).

By focusing on object rehearsal through location, our findings are also distinct from complementary recent studies that have linked directional biases in microsaccades to the rehearsal of visual shape and orientation information (56–60). In contrast to these studies where microsaccades directly tracked the target-relevant memory feature (shape or orientation), we used microsaccades to track incidental object-locations that were never queried for report.

Consistent with a role of the oculomotor system in object rehearsal, complementary dual-task experiments have revealed enhanced memory for objects whose incidental locations became saccade targets during the working-memory delay (20, 61). This suggests some type of “refreshing” of memory objects when they become the target of a saccade. Our microsaccade observations were made without asking participants to make any eye movements, and persisted for several seconds after encoding. Accordingly, our microsaccade observations may signal serial refreshing by the oculomotor system that may serve as a mechanism supporting object maintenance. In future studies, it will be interesting to address whether and how these microsaccade findings relate to recent complementary findings of serial object-sampling in working memory (e.g., 62–64).

### Why does the observed microsaccade-maintenance bias diminish with time?

Experiment 2 demonstrated that the directional microsaccade bias was specific to memory demands, ruling out lingering effects from passive and attentive encoding. At the same time, despite ongoing memory demands, the bias diminished with time in the delay interval, and this was the case in both experiments. Interestingly, however, this appears not to be unique to the working-memory signature that we uncovered here. For example, the Contralateral Delay Activity (an EEG signature frequently implicated in working-memory maintenance; e.g., 65–67), similarly decays during the memory delay (e.g., 68–70).

Even so, the observed decay raises the intriguing question of why this may occur and what it may reflect. One possible explanation is that rehearsal demands become less urgent as time passes. Early after encoding, active rehearsal may facilitate consolidation, while later in the delay active rehearsal may be less critical, and passive maintenance may suffice. For example, with time, working-memory representations may become increasingly reliant on activity-silent mechanisms (through temporary changes in synaptic connectivity, see e.g., 71, 72), rather than states of active rehearsal. Another possibility is that over time, location information simply becomes less relevant for working-memory maintenance, as objects are gradually recoded to a less visual format, or shifted from peripheral to more foveal (e.g., 73, 74) or global (75) cortical representations. Finally, it is conceivable that the anticipation of the probe stimulus itself may have affected our ability to measure robust microsaccade biases, since stimulus expectation is known to reduce overall microsaccade rates (76–79). It is worth noting that these explanations are not mutually exclusive, as multiple (as well as additional) factors may contribute jointly; providing a relevant avenue for future research.

### What opportunities and challenges do these findings bring?

Our findings demonstrate how microsaccades can be used to track working-memory rehearsal through spatial location over time. This may open new avenues for characterising the spatial-temporal trajectories of working-memory rehearsal in future studies with complementary manipulations of spatial and/or temporal working-memory demands. We have previously demonstrated that microsaccades are a valuable tool for studying attentional selection and prioritisation of objects in visual working memory (e.g., 42, 53, 54). Here, we extend this involvement to the maintenance period. Zooming out, these findings add to a growing body of work showing the utility of using microsaccades to track cognition at large, including for tracking complementary cognitive functions such as temporal expectations (76–79), perceptual learning (80) and decision making (81).

At the same time, our data bring a practical implication to the fore that presents a challenge, rather than an opportunity. They show how microsaccades cannot be neglected in neuroscience studies of visual working memory. Several recent reports already made clear how microsaccades may confound neural decoding of memory representations, by showing how microsaccades can be systematically biased by task-relevant object-features such as shape and orientation (56–60). Building on this work, our data make clear how systematic biases in microsaccades can be a concern also for arbitrarily chosen spatial stimulus-configurations at encoding, even if stimulus configurations are never explicitly relevant for the participant.

### Conclusion

In conclusion, by studying microsaccades, we provide unique evidence that the brain’s oculomotor system participates in rehearsing visual objects in working memory through their incidental locations. The microsaccade-maintenance bias we observed may signal an attentional refreshing mechanism by the oculomotor system that supports visual object maintenance. At the same time, the observed decay in the observed microsaccade-maintenance bias raises intriguing questions about the dynamic nature of working-memory rehearsal across time.

## METHODS

### Participants

We conducted two experiments with 25 healthy human volunteers in each (Experiment 1: mean age = 21.96 years, age range = 18-28 years, 13 women; 1 left-handed; Experiment 2, mean age = 21.76 years, age range = 18-37 years, 17 women; 1 left-handed). Sample size of 25 per experiment was determined a-priori based on the use of the same sample size in relevant prior studies from the lab that used the same outcome measure to target complementary questions (42, 53, 54, 82, 83). The recruitment of participants for the two experiments was conducted independently. Data from all participants were included in the analysis. All participants received course credit or monetary compensation (€10 per hour), provided written informed consent prior to participation, and reported normal or corrected-to-normal vision.

### Task and procedure Experiment 1

Experiment 1 used a within-subject design in which we manipulated stimulus configuration at encoding. Participants performed a visual working memory task (see **Fig. 1a**), in which they memorised the orientation and colour of two objects presented on opposite points around a central fixation dot. The key manipulation was the variation of the stimulus configuration at encoding (horizontal, vertical, or either of two diagonal configurations; see **Fig. 2a**, top). This manipulation enabled us to investigate whether microsaccades during the subsequent delay-interval were more closely aligned with the configurational axes of the memory content, as opposed to the orthogonal axes – even if object location was incidental to the task because a location was never asked about.

Each trial started with a one-second intertrial interval that featured a small fixation dot, followed by the simultaneous presentation of two memory objects for 200 ms and a delay-interval of 2800 ms. The colour of the fixation dot then changed for 1000 ms serving as a retrocue to instruct (with 100% validity) which visual memory object would be probed after the second delay interval (1000 ms). The probe was always tilted at an angle of ±20 degrees relative to the cued memory target. The probe was always black and remained on the center of the screen until a response was provided. Participants had to match the orientation of the probe with that of the cued memory object by pressing the F key (counterclockwise) or the J key (clockwise).

The experimental stimuli were generated and presented on an LCD monitor with a solid grey background (ASUS ROG Strix XG248; 1920 × 1080 pixels; 240 Hz refresh rate) using PsychToolbox (84) for MATLAB. Participants were seated in a dimly lit room with their heads resting on a chin rest at approximately 70 cm from the display. The central fixation dot was black and approximately 0.3 degrees visual angle. The memoranda had a diameter of 1.6 degrees, and were presented with a horizontal and vertical displacement of 6.4 degrees. The orientation of each stimulus was chosen randomly. The configuration (horizontal, vertical, and diagonal), colour (red and blue; 21/165/234; 234/74/21), and target position of the memoranda were varied pseudo-randomly to ensure that all these features were distributed equally within each block. The duration of each experimental session, which had 12 blocks of 36 trials each (totaling 432 trials), was roughly 70 minutes. Participants prepared for the experiment by practising 36 trials.

### Task and procedure Experiment 2

Experiment 2 was designed to determine whether the directional microsaccade bias observed in Experiment 1 reflected the active location-based rehearsal of memory content, or instead reflected a lingering trace from encoding. To this end, Experiment 2 followed the same overall set-up as Experiment 1 but introduced different task demands, that were blocked. Moreover, to reduce factors, in Experiment 2, we always presented objects in either of the two diagonal encoding configurations.

In the passive viewing blocks (**Fig. 1b**), participants were instructed to ignore the stimuli, while maintaining central fixation. In the perceptual-judgement blocks (**Fig. 1c**), participants had to indicate whether the orientations of the two objects that were concurrently presented in any given display were identical (∼7% of encoding displays) by pressing the spacebar within 1 second after stimulus onset. The task in the working-memory blocks (**Fig. 1d**) was the same as Experiment 1, only now the colour cue (informing the target memory object) was integrated into the probe by making the probe itself match the colour of either memory object. The orientation of each stimulus was chosen randomly, but, unlike Experiment 1, target and non-target objects always had a difference in orientation of at least 20 degrees (except in identical-target trials that served as targets in the perception task). This separation was necessary to account for the fact that, in perception blocks, participants had to be able to differentiate between targets (identical orientations) from non-target (non-identical orientations). Participants completed 24 blocks, each with 48 trials (totalling 1152 trials). The order of the blocks was randomised in sets of three (i.e. passive, perception, memory). Each block began with a brief overview of the instructions for the upcoming task. Participants practised 32 trials of each of the three task types in preparation for the experiment.

### Data acquisition & preprocessing

The right eye position was continuously recorded with an EyeLink 1000 Plus (SR Research, Ltd.) at a sampling rate of 1000 Hz. The eye-tracking data were converted into ASCII format and analysed with Matlab and Fieldtrip. Eye blinks were detected with an algorithm based on fluctuations in the pupil data (85) and linearly interpolated. The data were then segmented into epochs from -500 ms to 3000 ms relative to encoding onset. We estimated participants’ attentiveness to the memory task by measuring reaction times and excluding trials with responses slower than the mean reaction time +4 standard deviations (following an iterative procedure until no more outliers remained). We also did not analyse trials from perception blocks that contained targets (identical orientations) or were misidentified as such (false detection). Following these criteria, only a small percentage of trials were eliminated in each experiment (Experiment 1: *M* = 0.015%, SD = 0.008%; Experiment 2: *M* = 0.36%, *SD* = 0.008%).

### Eye-tracking analysis

To detect gaze shifts (saccades) we build on an existing velocity-based detection approach that has been successfully employed and validated in prior studies from our lab (42, 53, 86). Saccade detection was achieved by calculating the euclidean distance between successive gaze points in the horizontal and vertical planes, smoothing the resulting velocity vector (using the built-in “smoothdata” function in MATLAB, with a 7 ms sliding window), and identifying the onset of a saccade as the instant that the velocity exceeds a predetermined threshold. A trial-based threshold of 5 times the median velocity was employed to determine saccade onset, and a minimum interval of 100 milliseconds was used between saccades to prevent multiple detections of the same saccade.

To index the incidental use of locations for working memory, we tracked the direction (angle) of saccades and classified them as “aligned” when their direction was closer to the configurational axis of the visual objects during encoding, and “misaligned” when their direction was closer to the orthogonal axis. Every detected saccade was thus either classified as aligned or as misaligned. The resulting aligned and misaligned saccade-rate time courses (expressed in Hz) were then smoothed (using the built-in function “smoothdata” in MATLAB) using a moving average with a 100 ms sliding window. Crucially, directions classified as aligned in one configuration (e.g., vertical stimulus arrangement) would be classified as misaligned in the complementary configuration (e.g., horizontal stimulus arrangement). Accordingly, by collapsing across configurations, we were able to isolate the effect of stimulus configuration by subtracting out any background preferred saccade directions that would be common across all configurations.

### Statistical analyses

To assess whether the oculomotor system is involved in the rehearsal of visual objects in working memory, we compared the time courses of aligned and misaligned saccade rates. This was statistically evaluated using a cluster-based permutation approach (87), which effectively avoids the issue of multiple comparisons by evaluating the full dataspace under a single permutation distribution of the largest cluster. It groups neighbouring time points that show significant differences between the two time-courses into a cluster and then randomly permutes the condition labels to create a null distribution of the largest cluster that would be observed by chance. If the observed cluster size is larger than what would be observed by chance, the two time-courses differ statistically. We used the Fieldtrip toolbox (88) to identify significant clusters using the default configuration settings (10,000 permutations; alpha level of 0.025 per side).

In Experiment 2, we additionally compared the directional microsaccade biases from our working-memory condition of interest to each of the control conditions (passive, perception) using paired sample t-tests. For this comparison, we extracted the effect of interest as the difference in aligned vs. misaligned saccades rates, averaged across the full delay interval, as determined a-priori.

## Data availability

Raw data and relevant code will be made publicly available upon publication.

## ACKNOWLEDGEMENTS

This research was supported by an ERC Starting Grant from the European Research Council (MEMTICIPATION, 850636) and an NWO Vidi grant by the Dutch Research Council (grant number 14721) to F.v.E. The authors thank Anna van Harmelen and Baiwei Liu for their valuable comments on the manuscript.

